# Genome-wide polygenic score to identify a monogenic risk-equivalent for coronary disease

**DOI:** 10.1101/218388

**Authors:** Amit V. Khera, Mark Chaffin, Krishna G. Aragam, Connor A. Emdin, Derek Klarin, Mary E. Haas, Carolina Roselli, Pradeep Natarajan, Sekar Kathiresan

## Abstract

Identification of individuals at increased genetic risk for a complex disorder such as coronary disease can facilitate treatments or enhanced screening strategies. A rare monogenic mutation associated with increased cholesterol is present in ~1:250 carriers and confers an up to 4-fold increase in coronary risk when compared with non-carriers. Although individual common polymorphisms have modest predictive capacity, their cumulative impact can be aggregated into a polygenic score. Here, we develop a new, genome-wide polygenic score that aggregates information from 6.6 million common polymorphisms and show that this score can similarly identify individuals with a 4-fold increased risk for coronary disease. In >400,000 participants from UK Biobank, the score conforms to a normal distribution and those in the top 2.5% of the distribution are at 4-fold increased risk compared to the remaining 97.5%. Similar patterns are observed with genome-wide polygenic scores for two additional diseases – breast cancer and severe obesity.

**One Sentence Summary:** A genome-wide polygenic score identifies 2.5% of the population born with a 4-fold increased risk for coronary artery disease.

## Main Text

The identification of individuals at increased genetic risk for a common, complex disease can facilitate treatment or enhanced screening strategies to prevent disease manifestation. For example, with respect to coronary disease, ~1:250 individuals carry a rare, large-effect genetic mutation causal for increased low-density lipoprotein cholesterol (1-3). A recent analysis in a large U.S. health care system demonstrated that such individuals have an odds ratio for coronary disease of 2.6 when compared to non-carriers and an odds ratio of 3.7 for early-onset disease (1). Aggressive treatment to reduce circulating low-density lipoprotein cholesterol levels among carriers of such mutations can reduce coronary disease risk (4).

Beyond rare monogenic mutations, a decade of genome-wide association studies (GWAS) has demonstrated that common single nucleotide polymorphisms contribute to a range of complex diseases (5). However, because the effect size of such polymorphisms tends to be modest, any individual polymorphism has limited utility for risk prediction. Polygenic scores (PS) provide a mechanism for aggregating the cumulative impact of common polymorphisms by summing the number of risk variant alleles in each individual weighted by the impact of each allele on risk of disease (6). We recently demonstrated that a coronary disease PS consisting of 50 common variants that had achieved genome-wide levels of statistical significance in previous studies can stratify the population into varying trajectories of risk (7,8).

Simulated analyses based on GWAS effect size distributions suggest that the predictive power of such PSs may be markedly improved by considering a genome-wide set of common polymorphisms (9-11). But, it remains uncertain whether the extreme of a PS distribution can confer risk equivalent to a monogenic mutation (e.g., 4-fold increased risk). Here, we demonstrate that a PS comprised of a genome-wide set of common variants permits identification of individuals with 4-fold increased risk for coronary disease and subsequently generalize this approach to two additional complex diseases, breast cancer and severe obesity.

In order to develop an optimized polygenic score for coronary disease, we derived two new PSs and compared them with two previously published scores in a testing dataset of 120,286 individuals of European ancestry from the UK Biobank – 4,831 with coronary disease and 115,455 controls (7,12,13). The UK Biobank is a large observational study that enrolled individuals aged 45 to 69 years of age from across the United Kingdom beginning in 2006 (14).

We derived the two new PSs using summary association statistics from our earlier GWAS as a starting point for the relationship of millions of common polymorphisms to risk for coronary disease (Supp. Methods; 15). A reference population of 503 Europeans from the 1000 Genomes study was used to assess the correlation of a given polymorphism with others nearby (‘linkage disequlibrium’) (16). For the first score, we implemented a ‘pruning and thresholding’ strategy (PS_P&T_) to combine independent variants (r^2^ < 0.8 with other nearby variants) that exceeded nominal significance (p-value < 0.05) in the previous GWAS. For the second score, we used the recently developed LDPred computational algorithm (17). This involves a Bayesian approach to calculate a posterior mean effect for *all* variants based on a prior (effect size in the prior GWAS) and subsequent shrinkage based on linkage disequilibrium.

All four scores demonstrated robust association with coronary disease in the testing dataset. But, the newly-derived genome-wide polygenic score of 6.6 million common single nucleotide polymorphisms (PS_GW_) demonstrated the maximal area-under-the-curve of 0.64 and was selected for use in subsequent analyses (**Table 1**).

**Table 1.** Association of 4 polygenic scores with coronary disease in testing dataset of 120,286 individuals. Area-under-the curve and odds ratios determined via logistic regression adjusting for the first four principal components of ancestry. GWAS= genome-wide association study; SD= standard deviation; P&T= pruning and thresholding; GW= genome-wide

| Polygenic score | Derivation strategy | N<br>Variants | Area-under<br>the curve | Odds ratio<br>(per SD increment) |
| --- | --- | --- | --- | --- |
| Tada et al. (7) | Variants that had achieved genome-wide levels of statistical significance in prior GWAS ($p < 5 \times 10^{-8}$) | 50 | 0.59 | 1.38 |
| Abraham et al. (8) | Linkage-disequilibrium based thinning of variants from prior GWAS | 49,310 | 0.59 | 1.38 |
| PS<sub>P&T</sub> | Pruning based on statistical significance ($p < 0.05$) and linkage disequilibrium ($r^2 < 0.8$) of variants from prior GWAS | 116,859 | 0.62 | 1.54 |
| PS<sub>GW</sub> | **LDPred computational algorithm to assign weights to all available variants from prior GWAS via explicit modeling of linkage disequilibrium** | **6,630,150** | **0.64** | **1.67** |

Next, we sought to validate this score in an independent dataset of the remaining 288,890 individuals of European ancestry in the UK Biobank. Mean age was 57 years and 55% of the cohort was female. 8676 (3.0%) of the participants had been diagnosed with coronary disease, as defined based on verbal interview with a trained nurse or hospitalization for myocardial infarction or coronary revascularization in the electronic health record prior to enrollment.

We tested the hypothesis that individuals with high PS_GW_ might have risk equivalent to a monogenic coronary disease mutation (e.g., four-fold increased risk) by assessing progressively more extreme tails of the PS_GW_ distribution and comparing risk with the *remainder* of the population (**Table 2**; **Fig. 1A**). Across UK Biobank participants, PS_GW_ conformed to a normal distribution and individuals in the top 2.5% of the PS_GW_ distribution had a four-fold increased coronary disease risk (odds ratio 3.96) when compared with the remaining 97.5% of the population in a logistic regression model adjusted for age, sex, genotyping array, and the first four principal components of ancestry. We defined those individuals in the top 2.5% of the distribution as having high PS_GW_ in subsequent analyses.

**Fig. 1.**
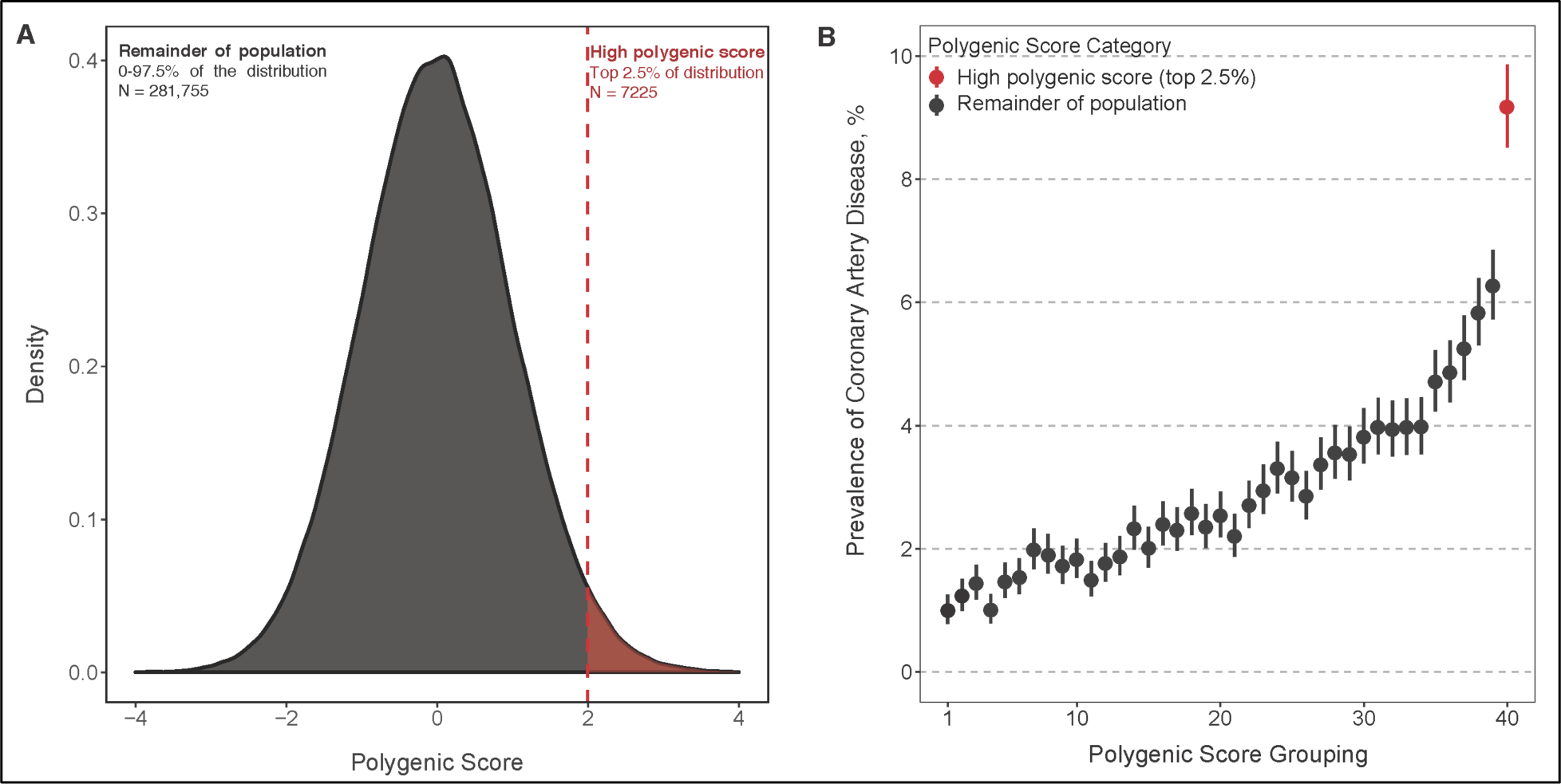
A new genome wide polygenic score (PS_GW_) identifies individuals with significantly increased risk of coronary disease. A near normal distribution of the PS_GW_ was noted in the UK Biobank validation cohort (**A**). The x-axis represents PS_GW_, with values scaled to a mean of 0 and standard deviation of 1 to facilitate interpretation. Individuals were binned into 40 groups based on PS_GW_, with each grouping representing 2.5% of the population (~7225 individuals). The high polygenic risk group displayed in red (top 2.5% of the distribution) had a significantly higher prevalence of coronary disease (**B**).

**Table 2.**
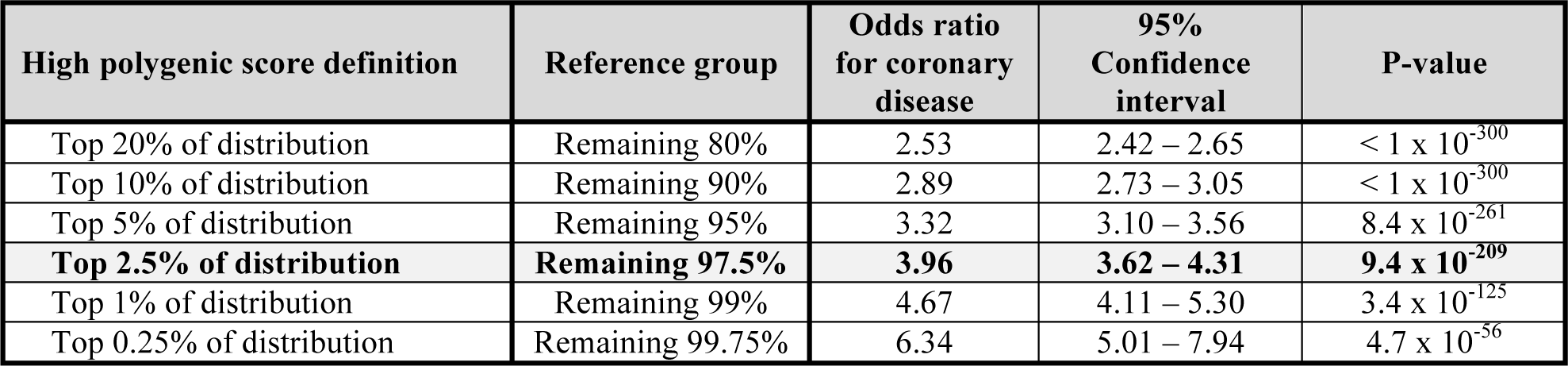
Prevalence and clinical impact of high polygenic score for coronary artery disease. Odds ratio for coronary disease calculated by comparing those with high polygenic score to the remainder of the population in a logistic regression model adjusted for age, sex, genotyping array, and the first four principal components of ancestry.

| High polygenic score definition | Reference group | Odds ratio for coronary disease | 95% Confidence interval | P-value |
| --- | --- | --- | --- | --- |
| Top 20% of distribution | Remaining 80% | 2.53 | 2.42 – 2.65 | $< 1 \times 10^{-300}$ |
| Top 10% of distribution | Remaining 90% | 2.89 | 2.73 – 3.05 | $< 1 \times 10^{-300}$ |
| Top 5% of distribution | Remaining 95% | 3.32 | 3.10 – 3.56 | $8.4 \times 10^{-261}$ |
| **Top 2.5% of distribution** | **Remaining 97.5%** | **3.96** | **3.62 – 4.31** | **$9.4 \times 10^{-209}$** |
| Top 1% of distribution | Remaining 99% | 4.67 | 4.11 – 5.30 | $3.4 \times 10^{-125}$ |
| Top 0.25% of distribution | Remaining 99.75% | 6.34 | 5.01 – 7.94 | $4.7 \times 10^{-56}$ |

Coronary disease was noted in 663 of 7225 (9.2%) individuals with high PS_GW_ as compared to 8013 of 281,755 (2.8%) of those in the remainder of the distribution (**Fig. 1B**). Of the 8676 individuals with coronary disease, 663 (7.6%) were predisposed on the basis of high PS_GW_. Several traditional coronary disease risk factors including family history of heart disease were enriched in those with high PS_GW_ (**Table 3**). However, attenuation in the risk estimate for high PS_GW_ was modest after additional adjustment for history of hypertension, type 2 diabetes, hypercholesterolemia, current smoking, and family history of heart disease (adjusted odds ratio 3.15; 95% confidence interval 2.86 – 3.46).

**Table 3.** Baseline characteristics according to high coronary disease polygenic score status. Values displayed are mean (standard deviation) for continuous variables and N (%) for categorical variables.

|  | **Remainder of population**<br>**(0 – 97.5% of distribution)** | **High polygenic score**<br>**(top 2.5% of distribution)** | **P-value** |
| --- | --- | --- | --- |
| Number of individuals | 281,755 | 7225 |  |
| Age, years | 56.9 (8.0) | 56.7 (8.1) | 0.01 |
| Male sex | 127,894 (45.4%) | 3189 (44.1%) | 0.04 |
| Hypertension | 78,999 (28.0%) | 2460 (34.0%) | < 0.001 |
| Type 2 diabetes | 13,547 (4.8%) | 441 (6.1%) | < 0.001 |
| Hypercholesterolemia | 38,001 (13.5%) | 1600 (22.1%) | < 0.001 |
| Current smoking | 25,908 (9.2%) | 691 (9.6%) | 0.29 |
| Family history of heart disease | 100,856 (35.8%) | 3364 (46.6%) | < 0.001 |
| Body mass index, kg/m<sup>2</sup> | 27.4 (4.7) | 27.7 (4.8) | < 0.001 |
| Systolic blood pressure, mmHg | 140 (19.7) | 141 (19.6) | < 0.001 |
| Lipid-lowering therapy | 47,550 (17.0%) | 1962 (27.3%) | < 0.001 |

In order to assess the generalizability of these observations, we used a similar approach to construct separate PSs for two additional complex diseases with major public health implications – breast cancer and severe obesity. As for coronary disease, we used summary association statistics from large prior GWASs as a starting point for the relationship of common polymorphisms to breast cancer or body-mass index (18,19).

Among 157,897 females of the UK Biobank validation dataset, 6567 (4.2%) had been diagnosed with breast cancer at the time of enrollment. Individuals with high PS for breast cancer had a 2.9-fold increased risk when compared with the remaining 97.5% of the population (**Table 4**). Breast cancer was noted in 10.5% of individuals with high PS as compared to 4.0% of those in the remainder of the distribution (Fig S1). Of individuals with breast cancer, 6.4% were predisposed on the basis of high PS. Attenuation in the risk estimate for high PS was modest after additional adjustment for family history of breast cancer, age at menarche, current smoking, body-mass index, and previous use of hormonal replacement therapy (adjusted odds ratio 2.78 95% confidence interval 2.49 – 3.09; Table S1)

**Table 4.**
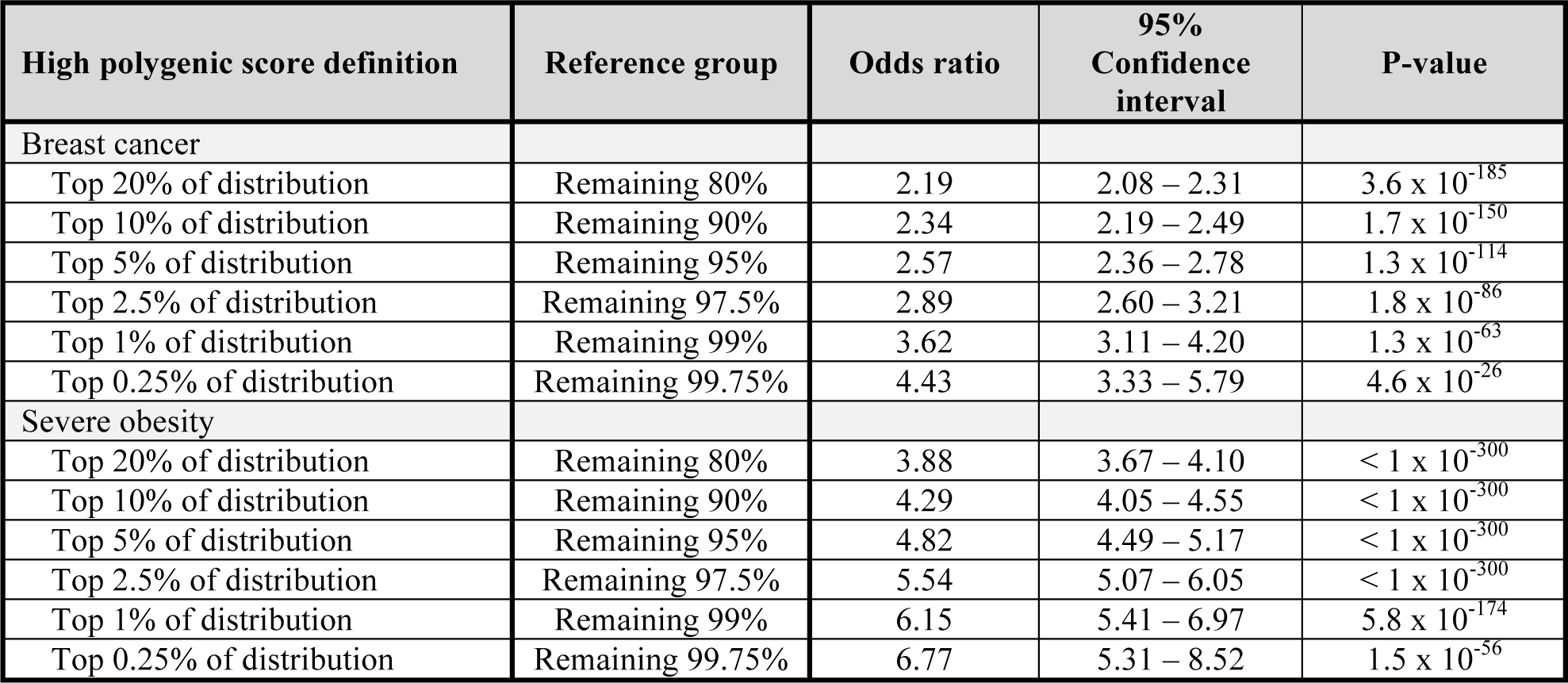
Prevalence and clinical impact of high polygenic score for breast cancer and severe obesity (body-mass index ≥ 40 kg/m^2^). Breast cancer analysis was restricted to females. Odds ratios calculated by comparing those with high polygenic score to the remainder of the population in a logistic regression model adjusted for age, sex (for severe obesity only), genotyping array, and the first four principal components of ancestry.

| High polygenic score definition | Reference group | Odds ratio | 95% Confidence interval | P-value |
| --- | --- | --- | --- | --- |
| Breast cancer |  |  |  |  |
| Top 20% of distribution | Remaining 80% | 2.19 | 2.08 – 2.31 | $3.6 \times 10^{-185}$ |
| Top 10% of distribution | Remaining 90% | 2.34 | 2.19 – 2.49 | $1.7 \times 10^{-150}$ |
| Top 5% of distribution | Remaining 95% | 2.57 | 2.36 – 2.78 | $1.3 \times 10^{-114}$ |
| Top 2.5% of distribution | Remaining 97.5% | 2.89 | 2.60 – 3.21 | $1.8 \times 10^{-86}$ |
| Top 1% of distribution | Remaining 99% | 3.62 | 3.11 – 4.20 | $1.3 \times 10^{-63}$ |
| Top 0.25% of distribution | Remaining 99.75% | 4.43 | 3.33 – 5.79 | $4.6 \times 10^{-26}$ |
| Severe obesity |  |  |  |  |
| Top 20% of distribution | Remaining 80% | 3.88 | 3.67 – 4.10 | $< 1 \times 10^{-300}$ |
| Top 10% of distribution | Remaining 90% | 4.29 | 4.05 – 4.55 | $< 1 \times 10^{-300}$ |
| Top 5% of distribution | Remaining 95% | 4.82 | 4.49 – 5.17 | $< 1 \times 10^{-300}$ |
| Top 2.5% of distribution | Remaining 97.5% | 5.54 | 5.07 – 6.05 | $< 1 \times 10^{-300}$ |
| Top 1% of distribution | Remaining 99% | 6.15 | 5.41 – 6.97 | $5.8 \times 10^{-174}$ |
| Top 0.25% of distribution | Remaining 99.75% | 6.77 | 5.31 – 8.52 | $1.5 \times 10^{-56}$ |

Among 288,018 individuals of the UK Biobank validation dataset with body-mass index available, 5232 (1.8%) were severely obese at the time of enrollment, defined as body-mass index ≥ 40 kg/m^2^. Individuals with high PS had a 5.5-fold increased risk of severe obesity when compared with the remaining 97.5% of the population (**Table 4**). Severe obesity was noted in 8.4% of individuals with high body-mass index PS as compared to 1.6% of those in the remainder of the distribution (Fig S2). Of individuals with severe obesity, 11.6% were predisposed on the basis of high PS. Results were similar when considering a less stringent definition for obesity of body-mass index ≥ 30 kg/m^2^ (Table S2).

For three common diseases, we demonstrate that the incorporation of a genome-wide set of common polymorphisms into a PS can identify subsets of the population at substantially increased risk.

These results permit several conclusions. First, we provide empiric evidence that the cumulative impact of common polymorphisms on risk of disease can approach that of rare, monogenic mutations. The predictive capacity of PSs will likely continue to improve as larger discovery GWAS studies more precisely define the effect sizes for common polymorphisms across the genome (9-11). Second, high PS_GW_ seems operable in a much larger fraction of the population as compared to rare monogenic mutations. For coronary disease, the largest gene-sequencing study to date identified a monogenic driver mutation related to increased low-density lipoprotein cholesterol in 94 of 12,298 (0.76%) afflicted individuals (1). Here, we identify high PS_GW_ in 7.6% of individuals with coronary disease, a prevalence an order of magnitude higher. Third, traditional risk factor differences of high PS_GW_ individuals versus the remainder of the distribution are modest and these individuals would thus be difficult to identify without direct genotyping. Fourth, a key advantage of a DNA-based diagnostic such as PS_GW_ is that it can be assessed from the time of birth, well before the discriminative capacity of most traditional risk factors emerges, and may thus facilitate intensive prevention efforts. For example, we recently demonstrated that high polygenic risk for coronary disease may be offset by adherence to a healthy lifestyle or cholesterol-lowering therapy with statin medications (8,20,21). Finally, we demonstrate similar patterns for two additional heritable diseases – breast cancer and severe obesity – suggesting that this approach will provide a generalizable framework for risk stratification across a range of common, complex diseases.

Several limitations deserve mention. First, the risk associated with a high polygenic score is not the result of a discrete underlying mechanism, but rather a quantitative blend of numerous risk pathways. Monogenic mutations predispose to disease on the basis of a specific driving pathophysiology that can sometimes enable targeted therapy. For example, homozygous deficiency in the *POMC* gene is associated with extreme obesity and precise targeting of the perturbed pathway can lead to significant weight loss (22). However, this monogenic etiology of obesity is exceedingly rare in the general population (23). To the extent that strategies to mitigate increased risk have utility regardless of underlying mechanism (e.g., statin therapy for coronary disease, dietary modification for severe obesity, or mammography screening for breast cancer), identification of individuals with high polygenic risk may prove useful. Second, the polygenic scores described here were derived and tested in individuals of European ancestry. Because allele frequencies, linkage disequilibrium patterns, and effect sizes of common polymorphisms vary by ancestry, future studies are needed to extend this approach across additional ancestral backgrounds (24). Lastly, the potential utility of genetic risk disclosure must be weighed against possible untoward consequences, including increased cost of care, psychological distress or discrimination, and a sense of fatalism in those at high risk. Additional work is needed to optimize genetic risk disclosure to patients and their health care providers and to test whether such disclosure can improve clinical outcomes.

## Acknowledgments

Dr. Khera reports funding support from a KL2/Catalyst Medical Research Investigator Training award from Harvard Catalyst funded by the National Institutes of Health (TR001100) and a Junior Faculty Research Award from the National Lipid Association. Dr. Natarajan received funding support from a John S. Ladue Memorial Fellowship from Harvard Medical School. Drs. Aragam and Klarin are supported by the National Heart, Lung, and Blood Institute of the National Institutes of Health (T32 HL007208 and T32 HL007734 respectively). Dr. Kathiresan is supported by an Ofer and Shelly Nemirovsky Research Scholar Award from Massachusetts General Hospital and RO1 HL127564 from the National Heart, Lung, and Blood Institute.

The authors thank Dr. David Altshuler (Vertex Pharmaceuticals; Boston, MA) for comments on an earlier version of this manuscript.

Analyses in the UK Biobank were conducted via application 7089 via a protocol approved by the Partners HealthCare Institutional Review Board.

## Supplementary Materials

Materials and Methods

Figures S1-S2

Tables S1-S2

